# Ethylene sensitivity assay for medicinal and fiber-type *Cannabis* seedlings reveals a triple-response-like phenotype

**DOI:** 10.1101/2024.08.28.610146

**Authors:** Adrian S. Monthony, Marilou Ledeuil, Davoud Torkamaneh

**Affiliations:** Département de phytologie, Université Laval, Québec City, Québec, Canada; Institut de Biologie Intégrative et des Systèmes (IBIS), Université Laval, Québec, Canada; Centre de recherche et d’innovation sur les végétaux (CRIV), Université Laval, Québec, Canada; Institut intelligence et données (IID), Université Laval, Québec, Canada; École d’ingénieurs en biotechnologies, Université Catholique de Lyon, Lyon, France

**Keywords:** *Cannabis sativa*, ethylene, ethephon, triple-response, silver thiosulfate, Arabidopsis, sexual plasticity

## Abstract

As evidence mounts for ethylene’s important role in sex determination, understanding ethylene signaling in *Cannabis sativa* L. (cannabis) has become increasingly vital. This study investigated the response of hemp-type and drug-type *Cannabis sativa* seedlings to ethylene, revealing a unique ‘paired response’ phenotype under dark conditions and ethephon treatment, suggesting a more specialized variation of the triple response distinct from that observed in other species. Employing a novel ethephon-based assay, this research bypassed the complexities of using gaseous ethylene, providing a more accessible method for examining ethylene responses. The results showed *C. sativa* seedlings exhibit marked sensitivity to ethephon at concentrations (125 mg/L, 250 mg/L, and 500 mg/L) lower than those previously reported to influence mature plants, indicating a broad ethylene responsiveness across various genetic backgrounds. Furthermore, the reversal of ethylene-induced phenotypic changes by silver thiosulfate (STS) at all concentrations tested (1, 2, and 3 mM) suggests the conservation of ethylene signaling pathways in *C. sativa*. Importantly these phenotypic responses were observed in both drug and hemp type cannabis. However, the absence of an exaggerated apical hook in treated seedlings potentially points to a unique regulatory mechanism in response to ethylene in achene-bearing plants. These findings, highlighted by the successful induction of ethylene-sensitive phenotypes, underscore the conserved nature of ethylene signaling in *C. sativa*. The results highlight the need for further investigations into the regulatory mechanisms of ethylene in cannabis, especially concerning sexual development, opening new pathways for optimized breeding and cultivation practices aimed at enhancing plant growth and reproductive strategies.

## Introduction

*Cannabis sativa* L. (cannabis), belongs to the Cannabaceae family and has been cultivated by humans for food, medicine and many material uses for thousands of years (Clarke, 1999). Over the centuries, this plant has undergone domestication and global expansion (Monthony, 2021). Today, much of the enthusiasm surrounding the plant is due to many of the purported medicinal benefits. These benefits stem from a group of secondary metabolites called cannabinoids, of which there are over 100 that have been identified including tetrahydrocannabinol (THC) and cannabidiol (CBD; Mudge *et al*., 2018, 2019). These cannabinoids have been studied for their medicinal benefits, with early research focusing on cannabinoids’ potential impact on conditions such as arthritis, anxiety, and autism (Blake *et al*., 2006; Nurmikko *et al*., 2007; Perucca, 2017; Barchel *et al*., 2019).

Cannabis is a diploid (2n=10) dioecious plant with an XX/XY sex chromosome pair (Monthony *et al*., 2024). It reproduces primarily by outcrossing via wind pollination, which has resulted in a genome that is highly heterozygous, despite many centuries of cultivation (Lapierre *et al*., 2023a). The genomic complexity has been further compounded by years of prohibition, leading to clandestine breeding programmes with poorly tracked pedigrees (Mudge *et al*., 2018). Due to these prohibitive circumstances, many essential tools relied upon by plant biologists remain undeveloped in *C. sativa* or have experienced significant delays in their development (Monthony *et al*., 2021c). For instance, methods for *in vitro* multiplication are only now beginning to emerge for large-scale plant production (Monthony *et al*., 2021a), and a replicable technique for somatic embryogenesis to regenerate transgenic plants *in vitro* is yet to be established (Monthony *et al*., 2021b). One class of techniques, common in well-studied plants like Arabidopsis but still unexplored in cannabis, consists of rapid assays designed to screen seeds for specific phenotypes (Guzman & Ecker, 1990). Such assays are characterized by ease of implementation, speed, and simplicity, rendering them indispensable to both plant breeding programs and studies involving mutant population screening. In the present study, a simple, low-tech, and rapid method for screening cannabis seedlings for ethylene-insensitive phenotypes, similar to the triple response assay in Arabidopsis, is introduced.

Ethylene is one of the most widely studied plant growth regulators, with a wide range of effects on plant biology including responses to abiotic and biotic stresses, fruit ripening and seed maturation as well as flower development and sex determination (Binder, 2020). In cannabis, the role of ethylene has recently come into focus as a regulator of sexual plasticity (Monthony *et al*., 2024). Sexual plasticity, initially characterized in the animal kingdom (Liu *et al*., 2017), manifests in plants as a phenomenon where a dioecious plant with sex chromosomes exhibits a phenotypic sex that contradicts its genotypic sex (Monthony *et al*., 2024). Ethylene plays a pivotal role in sex determination in numerous plants, notably within the monoecious Cucurbitaceae plant family (Manzano *et al*., 2014; Li *et al*., 2021). In cucurbits, it has been consistently observed that either exogenous treatments or elevated endogenous levels of ethylene tend to have feminizing properties, leading to an increased female-to-male flower ratio (Li *et al*., 2021). Conversely, inhibition of ethylene production or signalling has been reported to cause an increase in the male-to-female flower ratio (Boualem *et al*., 2015). The application of silver thiosulfate (STS), an inhibitor of endoplasmic reticulum-localized ethylene receptors (Binder & Schaller, 2017), has already been employed in cannabis breeding to generate male flowers with XX pollen from XX plants for self-pollination and the production of 100% XX (feminized) seeds (Lubell & Brand, 2018; Adal *et al*., 2021). Conversely, it has been demonstrated that the application of ethylene in the form of 2-chloroethylphosphonic acid (ethephon) to XY cannabis plants results in the induction of female flowers (IFFs; Ram & Jaiswal, 1970; Moon *et al*., 2020), pointing towards a reciprocal regulatory role for ethylene in sexual plasticity.

One approach to exploring this reciprocal ethylene regulation is to study it in mature, flowering cannabis plants, treated with ethylene or ethylene inhibitors (Mohan Ram & Sett, 1982b; Moon *et al*., 2020; Adal *et al*., 2021). However, raising a plant to maturity is a time-consuming process that can exceed four months, depending on the genotype, starting material and the desired stage of maturity for study (Rodriguez-Morrison *et al*., 2021; Lapierre *et al*., 2023b). An alternative system of study for ethylene response in cannabis are seedlings, which reduce the time required to identify plants with altered ethylene sensitivity, especially in the context of large-scale mutagenesis studies. Until now, a method for screening cannabis seedlings for altered ethylene sensitivity has not been published, which has limited the study of ethylene-related genes (ERGs) thus far.

The “triple response,” first observed by plant physiologist Dimitry Neljubow, involves three characteristic phenotypic alterations in etiolated seedlings grown in darkness in the presence of ethylene (Merchante & Stepanova, 2017). The phenotype is characterized by an increase in the diameter of the hypocotyl, inhibition of hypocotyl and root length growth and exaggeration of the curvature of the apical hook (Guzman & Ecker, 1990). This response remains largely conserved across dicots, some monocots, and can easily be distinguished visually (Merchante & Stepanova, 2017; Kieber & Schaller, 2019). Furthermore, seedling response to ethylene in model species *Arabidopsis thaliana* has been shown to be highly sensitive, even to minor deviations in a plants ability to biosynthesis or respond to ethylene (Merchante & Stepanova, 2017). Consequently, the triple response assay has become a favored tool for studying and identifying ERGs in Arabidopsis. The development of an assay which relies on similar physiological responses in *C. sativa* would represent a valuable and low-tech tool for advancing the study of ERG in the species.

The application of ethylene to seedlings is complicated by its gaseous nature, necessitating specialized air-tight environments and sensitive equipment to measure ethylene concentration (Zhang & Wen, 2010; Takahashi-Asami *et al*., 2016). An assay which uses an aqueous alternative to ethylene, such as ethephon, offers a simple way to study ethylene induced phenotypes in *C. sativa* seedlings. Ethephon has been shown to elicit ethylene response phenotypes in mature plants, with effective concentrations reported to vary significantly, ranging from approximately 500 mg/L to as high as 1920 mg/L (Ram & Jaiswal, 1970; Mohan Ram & Sett, 1982a,b; Moon *et al*., 2020). To date, there have been no studies on the effect of ethephon on *C. sativa* seedlings, although ethylene induced phenotypes have been reported from ethephon treated seedlings in a number of other species including maize and lupin (Ortuño *et al*., 1991; Liu *et al*., 2020). The present study demonstrates the use of ethephon to induce a triple response-like phenotype in *C. sativa* from both industrial/fiber-type (<0.3% THC) and medicinal/drug-type (>0.3% THC) phenotypes. By employing ethephon as an ethylene source, this research provides a rapid and accessible method for early investigating ethylene responses in *C. sativa* seedlings, facilitating understanding of ethylene’s role in plant development and sexual plasticity.

## Materials and Methods

### Preparation of ethephon and silver thiosulfate

Ethephon was prepared at a concentration of 500 mg/L, as outline by Moon *et al*. (2020) from a commercially available stock of 40% (AK Scientific, Union City, CA). The 500 mg/L (3.46 mM) solution was prepared in distilled water with 1% Tween 20 (Fisher Chemical, Fair Lawn, NJ), as a surfactant. The solution was stored in the dark at room temperature for a maximum of 1 month prior to replacement. Serial dilutions of 250 mg/L (1.73 mM) and 125 mg/L (0.87 mM) were prepared as needed development of the method. STS was prepared as outline in Jones & Monthony (2022). Briefly, 3 mM STS + 0.1% Tween 20 was prepared fresh before each application, by mixing a 1:1 ratio of sodium thiosulfate (24 mM; Fisher Chemical) stock and silver nitrate (6 mM; Fisher Chemical) stock. Stock solutions were prepared using distilled water, and stored for a maximum of 3 months.

### Optimal ethephon and STS concentrations

To determine the optimal concentration for inducing the triple response in *C. sativa*, a range of ethephon concentrations (0, 125, 250, and 500 mg/L) were tested (Table 1), and the seedling responses were quantified as outlined below. Each group of seeds (n=15) was immersed in 50 mL of the respective ethephon concentration for a 12-hour period. Following soaking, seeds were sown on humidified cotton within square Petri dishes. Square Petri dishes (Fisherbrand, Waltham, MA) were stood upright and filled halfway with cotton (Medline, Northfield, IL) and seeds were positioned on the top of the cotton. The cotton was moistened with the same ethephon solution concentration as the seeds prior to sowing. Before sealing the Petri dishes, the seeds were misted with distilled water using a spray bottle (approximately 15 sprays per Petri, regardless of the treatment). The Petri dishes were subsequently sealed with PVC sealing film (PhytoTech Labs, Lenexa, KS) and positioned upright in custom-made Petri dish holders (as shown in Figure 1; top left). They were stored in the dark for 7 days. Each Petri dish contained 5 seeds. Seeds used in this experiment were from the fiber-type hemp cv. Vega (Dedecker *et al*., 2020).

**Figure 1.**
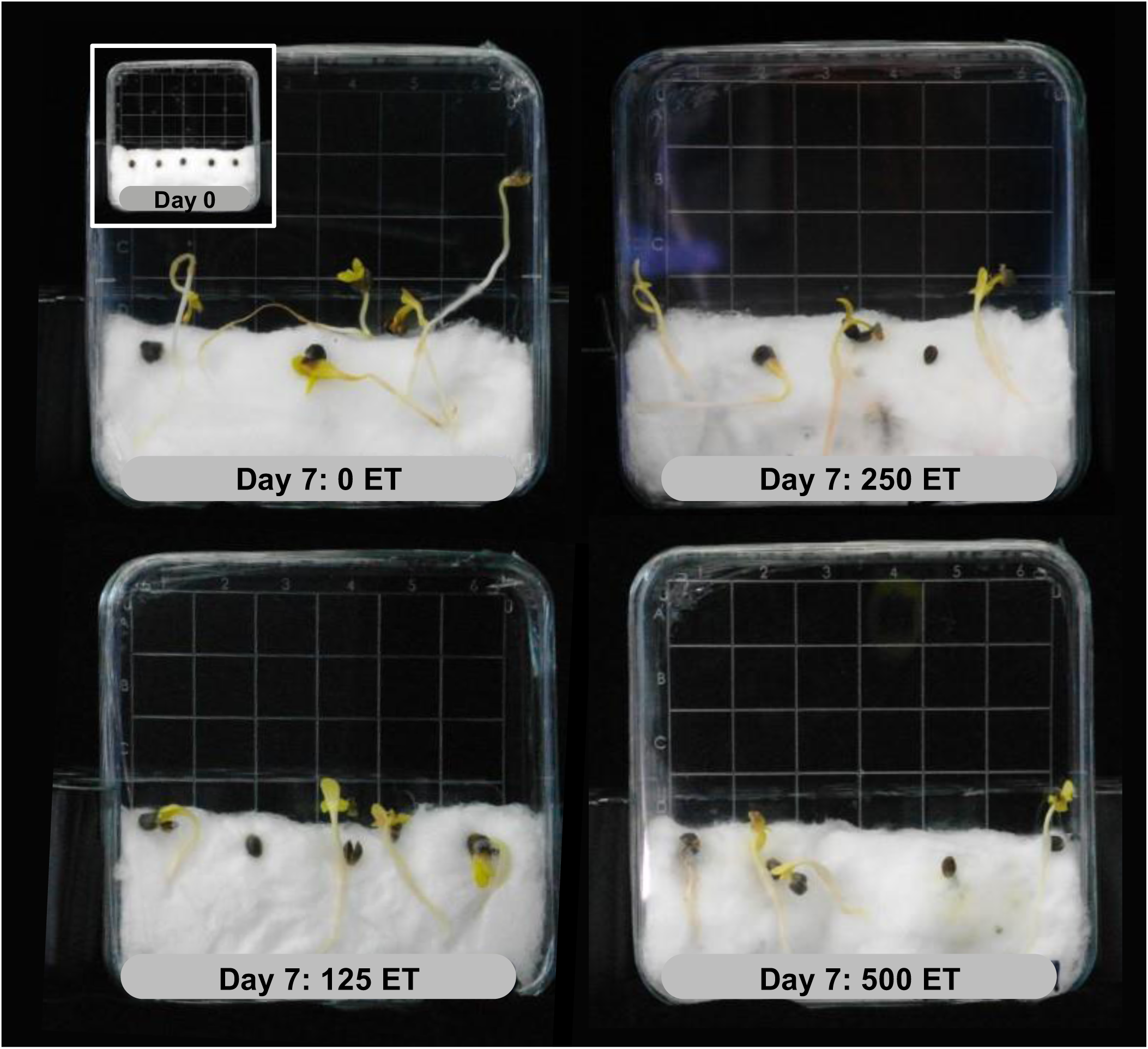
Morphological differences in *C. sativa* cv. Vega seedlings after 7 days of dark germination under varying ethephon (ET) concentrations. The tested concentrations (125, 250, and 500 mg/L) significantly affected the seedlings compared to the ethephon-free control (0 mg/L; 0 ET). The top left insert shows the initial layout of the seeds within the dish at day 0 for reference.

Once an optimal concentration of ethephon had been determined (250 mg/L), a second experiment was undertaken to determine the concentration of STS required to inhibit the altered seedling responses to ethephon. Five treatments were tested (Table 1). Petri dishes were set up using the same methodology. The key alteration made was the substitution of the misting solution, with the dishes being misted with various tested STS concentrations—specifically, 3 mM, 2 mM, and 1 mM—equivalent to approximately 15 sprays or about 6 to 7 mL per dish. Cotton was imbibed with 0 or 250 mg/L ethephon as indicated in Table 1.

**Table 1.** Experimental design and treatment optimization for the study involving *C. sativa* seedlings. Three experiments were conducted: (1) Optimization of ethephon concentration in hemp-type *C. sativa* to elicit a triple-response-like phenotype, (2) Optimization of STS (silver thiosulfate) concentration to restore the wild-type phenotype in hemp-type *C. sativa*, and (3) Testing the optimal concentrations of ethephon and STS, both individually and in combination, on drug-type *C. sativa*. The table summarizes the concentrations of ethephon and STS used in each experiment.

| 1. Optimisation of [ethephon] in hemp-type *C. sativa* |  | 2. Optimisation of [STS] in hemp-type *C. sativa* |  | 3. STS and ethephon alone, and in combination in drug-type *C. sativa* |  |
| --- | --- | --- | --- | --- | --- |
| [Ethephon]<br>(mg/L) | [STS]<br>(mM) | [Ethephon]<br>(mg/L) | [STS]<br>(mM) | [Ethephon]<br>(mg/L) | [STS]<br>(mM) |
| 0 | 0 | 0 | 0 | 0 | 0 |
| 125 | 0 | 250 | 0 | 0 | 1 |
| 250 | 0 | 250 | 1 | 250 | 0 |
| 500 | 0 | 250 | 2 | 250 | 1 |
|  |  | 250 | 3 |  |  |

### Triple response in drug-type *C. sativa*

Using the treatment optima determined in the prior experiments, the triple response assay was tested in a diverse population of feminized (XX) drug-type *C. sativa* seeds. Seeds were randomly sampled from a population of 79 different genotypes grown under conditions previously outlined in LaPierre *et al*. (2023b). Four different treatments were applied (Table 1), a total of 5 Petri dishes, each containing 5 seeds, were prepared per treatment (n=25) as already described.

### Data collection

At the end of the assay (day 7), morphological traits were assessed, including hypocotyl width, lengths, radicle length and apical hook angle. In order to obtain these measures, seedlings were individually photographed (Samsung NX100; Samsung Inc., San Jose, CA) and then analyzed using ImageJ (v1.54d, National Institute of Mental Health, Bethesda, MD) as described in Page *et al*. (2021). In brief once the scale was set, a straight-line tool was used to measure the width of the hypocotyls, and a freehand line was used to trace the lengths of the hypocotyl and radicles of each seedling. Apical hook angle measurements for each seed were obtained in triplicate for each sample, using the angle tool in ImageJ. All line tracing was done using a Wacom Intuos tablet (Wacom Co. Ltd., Saitama, Japan).

### Data analysis

All statistical analyses were conducted using R (v4.3.3) in the RStudio environment (v2024.04.2 Build 764, “Chocolate Cosmos” Release). A detailed description of the code used in these analyses, as well as the raw data, is available on the Open Science Framework (OSF) under the following DOI: 10.17605/OSF.IO/YBQ7F. The analyses were designed to evaluate the effects of ethephon and silver thiosulfate (STS), both independently and in combination, on traits associated with the triple response to ethylene in etiolated seedlings of drug- and hemp-type *Cannabis sativa*. The morphological traits analyzed included hypocotyl length, hypocotyl width, radicle length, and apical hook angle.

For each morphological trait, a Linear Mixed-Effects Model (LMM) was constructed using the ‘*lme’* function from the *nlme* package. Ethephon and STS were included as fixed factors in the models. Petri dishes were treated as random effects to account for variability between experimental units. Additionally, seeds nested within petri dishes were also treated as random effects to address the hierarchical structure of the data. In the case of apical hook angle, repeated measures of individual seeds nested within petri dishes were treated as random effects to account for the repeated observations and variability among experimental units. The assumptions of the LMMs were validated by examining the residuals for normality and homogeneity of variance. A custom function (‘ScaledResid’), used to scale residuals and correct for any covariance structures within the data, is available on the OSF page for this article. Normality of the model’s residuals was assessed visually using a Quantile-Quantile (QQ) plot and quantitatively using a Shapiro-Wilk test. Homogeneity of variance (homoscedasticity) across different levels of the fixed factors was evaluated using Levene’s test from the car package.

Depending on the outcome of these checks, data transformations, including rank transformations, were applied as necessary to meet model assumptions. The effectiveness of any transformation was re-evaluated using the same tests described above to ensure the transformed data met the model assumptions. Additionally, we compared the results of the untransformed and transformed models to assess the robustness of our models to potential deviations from normality. If the untransformed models demonstrated robustness and produced similar results to the transformed models, we proceeded to analyze the raw, untransformed data, favoring simplicity and interpretability.

Significant main effects and interactions identified by the LMMs were further explored using Tukey’s Honest Significant Difference (HSD) tests, performed via the emmeans package. These post-hoc analyses facilitated detailed pairwise comparisons between treatment groups. To visually represent the significance of differences between treatment groups, compact letter displays were generated using the multcompView package. Data visualization was performed using the ggplot2 package (Wickham, 2016), where compact letter displays were used to clearly depict the significance of differences among treatment groups.

## Results

All tested ethephon concentrations were observed to successfully induce the same triple response-like phenotype in the hemp-type *C. sativa* seedlings (Figure 1). This response was characterized by shortened hypocotyls and radicles, as well as increased hypocotyl width when compared to the control group, which did not receive ethephon. Remarkably, no significant differences were detected among the treatments, which included 500 mg/L, 250 mg/L, and 125 mg/L of ethephon (Table 2). When comparing the growth of untreated seeds to those treated with 250 mg/L of ethephon we found that the average hypocotyl and radicle lengths of untreated seeds were approximately double that of the ethephon-treated seedlings, as illustrated in Table 2. Additionally, the hypocotyl width of seedlings treated with 250 mg/L of ethephon doubled in comparison to untreated seedlings, measuring 2.22 cm as opposed to 1.15 cm. Notably, there was no exaggeration of the apical hook in any of the ethephon treatments compared to the control (Table 2). A concentration of 250 mg/L was selected for subsequent experiments, as it appeared effective in inducing a triple response like phenotype in dark grown seedlings.

**Table 2.** Effect of ethephon concentration on hemp cv. Vega dark-germinated seedlings. Estimated marginal means (emmeans) within the same column, followed by different letters, are significantly different according to a Tukey-Kramer HSD post-hoc test conducted following the Linear Mixed-Effects Model (LMM) analysis (α = 0.05). The number of germinated seeds measured (*n*) is reported out of the total of 15 seeds in each treatment.

| [Ethephon]<br>(mg/L) | [STS]<br>(mM) | $n$ | Hypocotyl Length<br>(mm) | | | | Hypocotyl Width<br>(mm) | | | | Radicle Length<br>(mm) | | | | Apical Hook<br>Angle (°) | | | |
| --- | --- | --- | --- | --- | --- | --- | --- | --- | --- | --- | --- | --- | --- | --- | --- | --- | --- | --- |
| 0 | 0 | 13 | 20.1 | ± | 2.3 | a | 1.2 | ± | 0.2 | a | 38.4 | ± | 6.1 | a | 148.3 | ± | 21.7 | a |
| 125 | 0 | 9 | 10.7 | ± | 1.2 | b | 2.2 | ± | 0.2 | b | 17.8 | ± | 1.3 | b | 107.9 | ± | 25.5 | a |
| 250 | 0 | 12 | 10.5 | ± | 1.0 | b | 2.2 | ± | 0.2 | b | 23.2 | ± | 2.1 | b | 104.5 | ± | 22.5 | a |
| 500 | 0 | 9 | 10.3 | ± | 1.2 | b | 2.3 | ± | 0.2 | b | 20.2 | ± | 1.8 | b | 111.5 | ± | 25.5 | a |

When tested in combination with different levels of STS, the effect of 250 mg/L ethephon was effectively inhibited by the addition of STS. All concentrations of STS produced similar inhibition of the triple response-like phenotype; the resulting phenotypes were indistinguishable from the untreated etiolated seedlings (Table 3). No changes in apical hook angle were observed in any of the treatments (Table 3). Among the tested STS concentrations, 1 µM of STS was selected for further evaluation in drug-type *C. sativa* seeds, as it quantitatively and qualitatively produced seedlings most closely resembling the control (untreated) seedlings.

**Table 3.** Effect of STS concentration in combination with ethephon on hemp cv. Vega dark-germinated seedlings. Estimated marginal means (emmeans) within the same column, followed by different letters, are significantly different according to a Tukey-Kramer HSD post-hoc test conducted following the LMM analysis (α = 0.05). The number of germinated seeds measured (*n*) is reported out of the total of 15 seeds in each treatment.

| [Ethephon]<br>(mg/L) | [STS]<br>(mM) | $n$ | Hypocotyl Length<br>(mm) | | | Hypocotyl Width<br>(mm) | | | Radicle Length<br>(mm) | | | Apical Hook<br>Angle ($^{\circ}$) | | |
| --- | --- | --- | --- | --- | --- | --- | --- | --- | --- | --- | --- | --- | --- | --- |
| 0 | 0 | 12 | 30.5 | ± 4.4 | b | 1.1 | ± 0.1 | a | 44.9 | ± 6.2 | b | 115.7 | ± 14.5 | a |
| 250 | 0 | 9 | 7.6 | ± 3.9 | a | 2.1 | ± 0.1 | b | 18.2 | ± 4.0 | a | 142.4 | ± 16.3 | a |
| 250 | 1 | 13 | 31.1 | ± 5.6 | b | 1.2 | ± 0.1 | a | 52.5 | ± 8.0 | b | 128.3 | ± 12.8 | a |
| 250 | 2 | 13 | 34.9 | ± 5.5 | b | 1.1 | ± 0.1 | a | 64.3 | ± 8.1 | b | 127.0 | ± 12.8 | a |
| 250 | 3 | 11 | 25.3 | ± 5.4 | ab | 1.2 | ± 0.1 | a | 44.6 | ± 6.5 | b | 145.3 | ± 13.9 | a |

Lastly the effects of STS alone and in combination with ethephon on drug-type *C. sativa* seedlings were explored. When STS was applied alone, it did not significantly affect the etiolation of the seedlings grown in the dark, as demonstrated in Table 4 and Figure 2. However, treating the dark-germinated seedlings with 250 mg/L of ethephon was found to be sufficient to induce a triple-response-like phenotype. This resulted in a significant reduction in hypocotyl and radicle lengths, along with a notable thickening of the hypocotyl, as shown in Figure 2. Importantly, the application of STS in combination with ethephon in drug-type *C. sativa* seedlings was identical to the response reported in the hemp-type counterpart, restoring the etiolated phenotype. No changes to apical hook angle were observed, as observed in the hemp-type *C. sativa* seedlings (Table 4).

**Figure 2.**
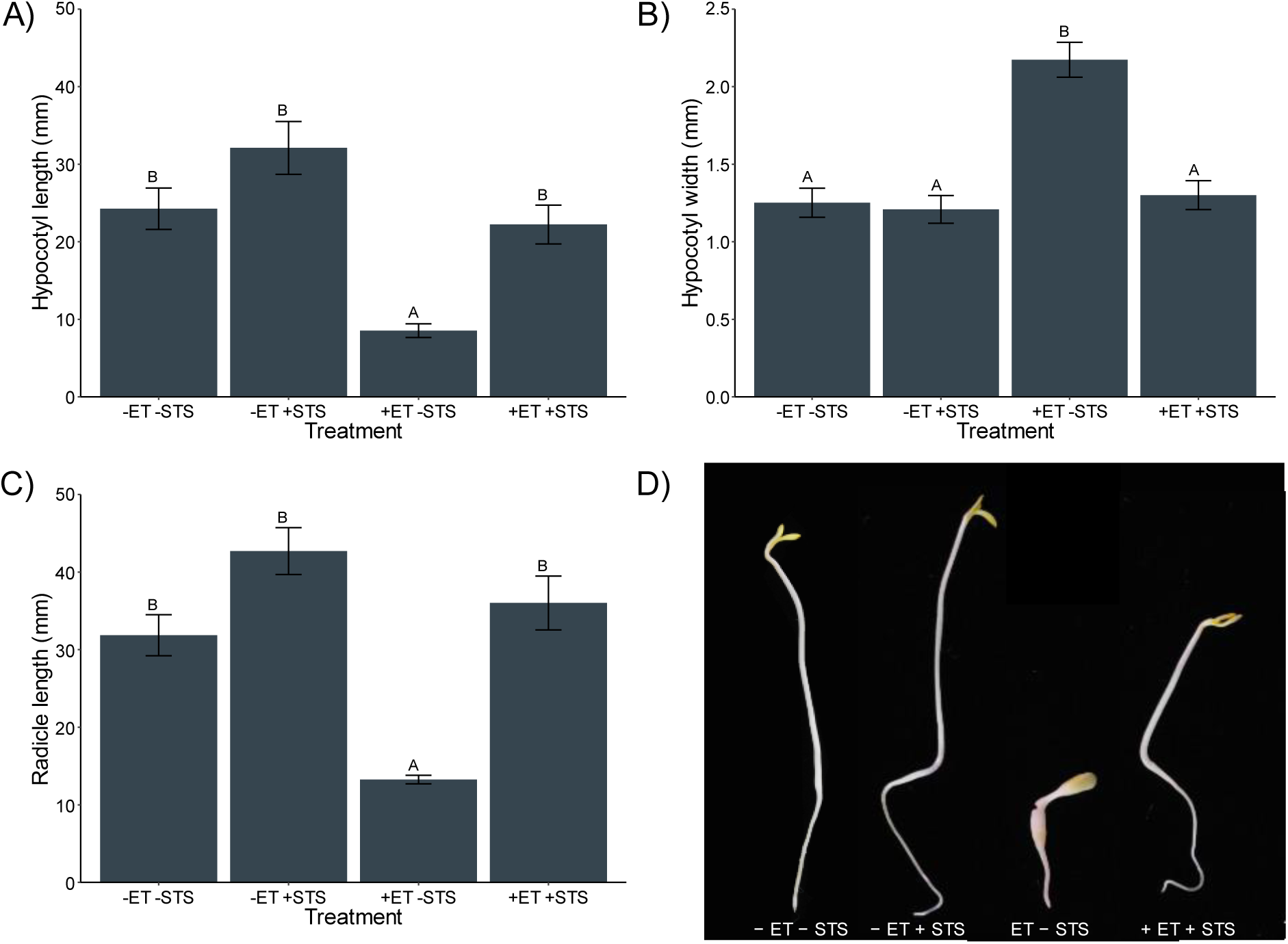
Triple response in drug-type *C. sativa.* A) Hypocotyl length; B) Hypocotyl width; C) Radicle length; D) Representative photos of seedlings grown in the presence (+) and/or absence (−) of ethephon (ET) and silver thiosulfate (STS). Error bars represent standard error of the mean. Scale bar = 5 mm.

**Table 4.** Assessment of the triple response in a feminized, drug-type *Cannabis sativa* mixed seed population. Estimated marginal means (emmeans) within the same column, followed by different letters, are significantly different according to a Tukey-Kramer HSD post-hoc test conducted following the LMM analysis (α = 0.05). The number of germinated seeds measured (*n*) is reported out of the total of 25 seeds used in each treatment.

| [Ethephon]<br>(mg/L) | [STS]<br>(mM) | $n$ | Hypocotyl Length<br>(mm) | | | Hypocotyl Width<br>(mm) | | | Radicle Length<br>(mm) | | | Apical Hook<br>Angle (°) | | |
| --- | --- | --- | --- | --- | --- | --- | --- | --- | --- | --- | --- | --- | --- | --- |
| 0 | 0 | 11 | 24.3 | ± 2.7 | b | 1.3 | ± 0.1 | a | 31.9 | ± 2.6 | b | 130.9 | ± 7.4 | a |
| 0 | 1 | 7 | 32.1 | ± 3.4 | b | 1.2 | ± 0.1 | a | 42.7 | ± 3.0 | b | 135.2 | ± 6.8 | a |
| 250 | 0 | 11 | 8.5 | ± 0.9 | a | 2.2 | ± 0.1 | b | 13.2 | ± 0.5 | a | 129.2 | ± 9.3 | a |
| 250 | 1 | 13 | 22.2 | ± 2.5 | b | 1.3 | ± 0.1 | a | 36.0 | ± 3.5 | b | 137.7 | ± 7.4 | a |

## Discussion

In this study, we explored the effects of ethylene, applied as ethephon, on dark-grown etiolated *Cannabis sativa* seedlings, focusing on the physical responses of the hypocotyl and radicle, and the apical hook formation, as has been previously characterized in Arabidopsis (Guzman & Ecker, 1990). *Cannabis sativa* exhibits a broad spectrum of sex expression, ranging from sexually dimorphic dioecious plants in medicinal strains to both monomorphic and dimorphic types in industrial hemp (Spitzer-Rimon *et al*., 2019; Dowling *et al*., 2024; Shi *et al*., 2024). Additionally, dioecious genotypes display sexual plasticity, capable of producing reproductive organs contrary to their chromosomal sex (Monthony *et al*., 2024). This sexual plasticity has been linked to the modulation of ethylene levels in both medicinal and industrial hemp-type *C. sativa* (Lubell & Brand, 2018; Monthony *et al*., 2024), but the molecular signalling mechanisms controlling this have not yet been identified in cannabis. There are several ongoing efforts to develop effective transformation methods for cannabis *in vitro* as well as *ex vitro* (Schachtsiek et al., 2019; Galán-Ávila et al., 2021; Ajdanian et al., 2023). These up-and-coming biotechnological tools are of interest in studying ERGs, which may provide a viable target for creating sex-stable cultivars of cannabis, thereby mitigating risks associated with sex changes in commercial cultivation. To help elucidate the molecular mechanisms controlling sexual plasticity and to screen for mutants with altered ethylene sensitivity, a rapid and straightforward test for ethylene sensitivity in seedlings is needed.

Methods for testing ethylene sensitivity have been developed in etiolated seedlings of other plant species such as *Arabidopsis thaliana* (Guzman & Ecker, 1990), pea (Beyer, 1976) and lupin (Ortuño *et al*., 1991). In these species, exposure of dark-grown, etiolated seedlings to ethylene results in the triple response, a phenotype characterized by shortening of the hypocotyl and radicle, swelling of the hypocotyl, and an exaggeration of the apical hook (Guzman & Ecker, 1990). In contrast, our study reveals a paired response in *C. sativa* seedlings treated with ethephon, characterized by the shortening of the hypocotyl and radicle, coupled with a swelling of the hypocotyl but without the exaggerated apical hook. This variation indicates that while ethylene signaling pathways may be largely conserved in *C. sativa*, there may be unique regulatory mechanisms, as suggested by the absence of the apical hook exaggeration.

The molecular mechanism underpinning ethylene’s influence on hypocotyl elongation has been clarified in *Arabidopsis* (Zhong *et al*., 2012). Notably, ethylene’s effect on hypocotyl growth is reversible and dependent on light conditions: it inhibits growth in the absence of light or under low-light conditions (<10 µmol m^-2^ s^-1^) and promotes hypocotyl elongation well-lit environments (Yu & Huang, 2017). This dual action of ethylene is mediated through the action of ETHYLENE INSENSITIVE 3 (EIN3) and ETHYLENE RESPONSE FACTOR 1 (ERF1), with ethylene exposure in dark conditions leading to an increase in EIN3 levels, which in turn promotes the activity of the ERF1 transcription factor (Zhong *et al*., 2012). This activation initiates the pathway that inhibits hypocotyl growth (Zhong *et al*., 2012). Our study demonstrates a significant reduction in hypocotyl elongation across all ethephon concentrations (Table 2), suggesting the EIN3 and ERF1 mediated growth inhibition mechanism could be conserved in *C. sativa*. Interestingly, no significant difference in hypocotyl length was observed between the highest and lowest ethephon concentrations, potentially due to a receptor saturation effect occurring at a concentration between 0 and 125 mg/L (Table 2).

This study investigates the effects of ethephon on cannabis seedlings, highlighting a significant gap in the literature regarding optimal ethephon concentrations for biological responses in this species, which has primarily been explored in mature plants (Mohan Ram & Sett, 1982a,b; Moon *et al*., 2020). Our findings reveal that *C. sativa* seedlings demonstrate a heightened sensitivity to ethephon at concentrations much lower than those known to affect mature plants, without noticeable variation across these reduced levels. This result contrasts sharply with previous research on mature *C. sativa*, which responded to a higher concentration range-about 500 mg/L to 1920 mg/L-to trigger physiological responses (Ram & Jaiswal, 1970; Mohan Ram & Sett, 1982a,b; Moon *et al*., 2020). Our data show that seedlings are receptive to much lower concentrations, with as little as 125 mg/L prompting a physiological ethylene response (Table 2). Interestingly, seedlings treated with 500 mg/L in our study showed similar responses to those observed at lower doses, indicating that seedlings have a broad spectrum of sensitivity to ethephon without showing signs of phytotoxicity at the highest level tested. The range of concentrations to which seedlings respond starts at much lower levels and may extend up to concentrations effective in mature plants. For example, concentrations like 250 mg/L, which were previously deemed ineffective in mature specimens (Ram & Jaiswal, 1970), elicited a paired response similar to that found at other tested concentrations in our seedlings (Table 3 and Figure 2). When comparing our results to those from seedlings of other species, it becomes evident that *C. sativa* seedlings share a heightened sensitivity to ethephon. For instance, *Lupinus albus L.* seedlings showed inhibited hypocotyl growth across ethephon concentrations from 0.66 mM (∼95 mg/L) to 6.6 mM (∼954 mg/L), with even the lowest concentration resulting in a transient growth reduction (Ortuño *et al*., 1991). Similarly, monocotyledonous plant seedlings, such as maize, exhibited stunted growth and increased mesocotyl coarseness in response to ethephon ranging from 0.001 mM (∼0.15 mg/L) to 1.2 mM (∼173 mg/L), in a dose-dependent manner (Liu et al., 2020). These findings are consistent with our observations in *C. sativa* seedlings, where all tested ethephon concentrations, starting from 125 mg/L, led to growth reduction over a 7-day period. This reinforces the idea that while mature plants may not be as sensitive to lower concentrations of ethephon, seedlings, including those of *C. sativa*, are highly responsive, suggesting that lower concentrations can effectively elicit physiological responses during early developmental stages.

In our study, the ethylene-induced phenotypic changes in *C. sativa* seedlings were completely reversed by the application of silver thiosulfate (STS), a known inhibitor of ethylene signaling (Figure 2, Table 3 Table 4). The silver ions (Ag^+^) in STS bind to the copper ion binding sites on ethylene receptor proteins in the endoplasmic reticulum, resulting in an inhibition of the receptor’s ability to bind ethylene and preventing the downstream ethylene-responsive gene expression (McDaniel & Binder, 2012; Binder, 2020). The reversal of the ethylene-induced phenotype by STS further suggests a conserved ethylene signalling mechanism in *C. sativa*, as has recently been proposed (Monthony *et al*., 2024). However, to fully validate this hypothesis, further transcriptomic and metabolomic analyses across multiple *C. sativa* cultivars are recommended, especially taking into account the species’ documented variability in *in vitro* responses to other plant growth regulators (Page *et al*., 2021; Monthony *et al*., 2021b).

The formation of an apical hook in germinating seedlings is crucial for protecting the meristem during soil emergence, and its development is known to be influenced by factors such as mechanical pressure, ethylene, and auxin gradients, as demonstrated in *Arabidopsis* (Shen *et al*., 2016). However, the exaggerated apical hook commonly seen as part of the ethylene-induced triple response is not universal across species. In achene-producing plants, light exposure, rather than ethylene, may play a more significant role in apical hook exaggeration, facilitating seed coat shedding during soil emergence (Shichijo *et al*., 2010). Given *C. sativa*’s classification as an achene-bearing plant (Jones & Monthony, 2022), this may explain the absence of an exaggerated apical hook in ethephon-treated seedlings in our study, despite significant reductions in hypocotyl length and increases in hypocotyl thickness.

Interestingly, this absence of apical hook exaggeration in *C. sativa* seedlings mirrors findings in other species, such as tomato and maize, where ethylene’s influence on the apical hook is minimal or reversed under certain conditions (Takahashi-Asami *et al*., 2016; Liu *et al*., 2020). These differences across species underscore the species-specific nature of ethylene responses in seedling development. While the exact reasons for the lack of an exaggerated apical hook in *C. sativa* remain unclear, it is possible that prolonged or more intense ethephon treatments could yield different outcomes, or that *C. sativa* exhibits a “paired response” similar to those seen in tomato and maize. Overall, the ability to reproduce a paired response to ethylene treatment in hemp and drug-type seedlings represents an important step toward understanding the complex role of ethylene in *C. sativa*, and establishes an effective and easily distinguishable method for future studies on ethylene signaling in cannabis.

## Conclusion

The exploration of ERGs in *C. sativa* is of growing importance as researchers begin to probe the role of ethylene in cannabis sexual plasticity. This study demonstrates for the first time that dark-germinated *C. sativa* seedlings show a triple response-like phenotype, which we have termed a “paired response”, when in the presence of ethylene. We have successfully introduced an ethephon-based assay that obviates the necessity for ethylene gas, facilitating a more accessible approach for researchers, especially those lacking specialized equipment to handle ethylene gas directly. Importantly, our findings indicate a broad ethylene sensitivity among etiolated *C. sativa* seedlings across diverse genetic backgrounds, including both hemp-type and drug-type genetic backgrounds. The effective reversal of ethylene-induced phenotypes by silver thiosulfate (STS) further supports the notion of the canonical ethylene signaling pathway in *C. sativa*, which we have proposed previously (Monthony & Jones, 2024). Nevertheless, the lack of an exaggerated apical hook in response to ethephon treatment in *C. sativa* seedlings hints at distinctive regulatory mechanisms possibly inherent to achene-bearing plants. This intriguing aspect merits further investigative efforts to unravel. The establishment of this assay offers a novel and valuable tool for the study of ethylene in cannabis, enriching the existing, yet limited, methodological arsenal. Particularly, this assay holds promise for future investigations into the ethylene-mediated control of sexual plasticity in cannabis. Understanding the dynamics of sexual plasticity will have profound implications for both the cultivation and breeding of *C. sativa*, enabling more precise manipulation of plant development and reproductive strategies. The results presented in this study contribute to our understanding of ethylene’s role in cannabis, in addition to paving the way for innovative research into the complex interplay of hormonal signaling and sexual development in the species.

## Acknowledgments

The authors gratefully acknowledge the support of the Natural Sciences and Engineering Research Council (NSERC) of Canada. ASM has also been supported by a NSERC Canada Vanier Graduate Scholarship. ML research internship was generously supported by the Université Catholique de Lyon École d’ingénieurs en biotechnologies as well as by the Bourse région mobilité intern from the Région Auvergne-Rhône-Alpes.

## Author Contributions

ASM, and DT planned and designed the research. ASM and ML performed the investigation and contributed to data curation. ASM and ML conducted formal analysis. ASM and DT acquired funding for the project. ASM and ML contributed to the methodology. ASM handled project administration. DT provided resources. ASM and DT supervised the research. ASM and ML visualized the results. ASM and ML wrote the original draft of the manuscript. ASM, ML, and DT contributed to reviewing and editing the manuscript.

## Data Availability Statement

All scripts and raw data used in this study are available from The Open Science Framework (OSF) under the DOI:10.17605/OSF.IO/YBQ7F and at https://osf.io/ybq7f/.

